## Supplemental Tables and Figures for "Low Gut Microbial Diversity Augments Estrogen-driven Pulmonary Fibrosis in Female-Predominant Interstitial Lung Disease"

**Table S1.** Mouse strains used in this study

| **Mouse Strain** | **Description** | **Designated Name** | **Vendor** |
| --- | --- | --- | --- |
| Wild-type (WT) C57BL/6J | C57BL/6 is the most widely used inbred strain. It is commonly used as a general purpose strain and background strain for the generation of congenics carrying both spontaneous and induced mutations. | WT | Jackson Laboratory  (Stock No: 000664) |
| Estrogen receptor alpha knock out mice (ERα KO mice) | B6DN (cg) – Esr 1<tm4.2ksk>/J  These mice carry a knock out mutation of the Esr1 gene in which exon 3 has been deleted. Homozygotes are viable but not fertile. | Esr-1^-/-^ | Jackson Laboratory  (Stock No: 026176) |
| Wild-type (WT) C57BL/6J Sham operated mice | Sham Surgery (faked surgical intervention that omits the ovariectomy) | WT Sham | Jackson Laboratory  (Stock No: 000664) |
| Wild-type (WT) C57BL/6J Ovariectomized mice | Ovariectomy (Female mice had their ovaries surgically excised). | OVX | Jackson Laboratory  (Stock No: 000664) |

**Table S2.** Flow cytometric antibodies used in this study

| **Antibodies against human proteins** | | | | |
| --- | --- | --- | --- | --- |
| **Antibody** | **Vendor, Catalog No** | **Host organism; antibody type** | **Clone** | **Conc (µg)/volume (µL) per test*** |
| CD3 Alexa Fluor 700 | BD 557943 | Mouse Monoclonal | UCHT1 | 1.25 µl |
| CD3 BV785 | Biolegend 317330 | Mouse Monoclonal | OKT-3 | 5 µl |
| IL-6 APC | BD 561441 | Rat Monoclonal | MQ2-13A5 | 5 µl |
| IL-17A PE | Biolegend 512306 | Mouse Monoclonal | BL168 | 5 µl |
| STAT3 (pY705) Pacific blue | BD 560312 | Mouse Monoclonal | 4/P-STAT3 | 10 µl |
| Lap TGF-β1 PerCP/Cy 5.5 | Biolegend 349612 | Mouse Monoclonal | TW4-2F8 | 1.6 µl |
| Lap TGF-β1 APC | Biolegend 349706 | Mouse Monoclonal | TGW4-6H10 | 1.6 µl |
| RORC AF488 | BD 563621 | Mouse Monoclonal | Q21-559 | 2.5 µl |
| **Antibodies against murine proteins** | | | | |
| CD3 Alexa Fluor 700 | Biolegend 100216 | Rat Monoclonal | 17A2 | 0.25 µg |
| CD4 APC-Cy7 | Biolegend 100414 | Mouse Monoclonal | GK1.5 | 0.1 µg |
| IL-17A PE | Biolegend 506904 | Rat Monoclonal | TCII-18H10.1 | 0.1 µg |
| IL-17A BV 650 | Biolegend 506930 | Rat Monoclonal | TCII-18H10.1 | 1.6 µg |
| STAT3 (pY703) Pacific Blue | BD 560312 | Mouse Monoclonal | 4/P-STAT3 | 10 µl |
| IL-6 FITC | Invitrogen 11-7061-82 | Rat Monoclonal | MP5-20F3 | 1.6 µg |
| IL-6 APC | Biolegend 504808 | Rat Monoclonal | MP5-20F3 | 0.04 µg |
| IL-23R APC | Biolegend 150906 | Rat Monoclonal | 12B2B64 | 0.02 µg |
| GP-130 PE | Biolegend 149404 | Rat Monoclonal | 4H1B35 | 0.02 µg |
| TGF-β1 PerCP/Cy5.5 | Biolegend 141410 | Mouse Monoclonal | TW7-16B4 | 0.13 µg |
| LIVE DEAD | Invitrogen L34967 |  |  | 1:1000 |

*Test = 1 million cells in 50µl of staining buffer

**Table S3.** Genomic regions demonstrating ERα binding in T-cells in the locus of *STAT3*. Chromosome positions are given in hg38 coordinates. Statistics are calculated by MACS2.

| **Chromosome** | **Start** | **End** | **Peak length** | **-log10(p-value )** | **Fold enrichment over input** | **-log10(q-value)** |
| --- | --- | --- | --- | --- | --- | --- |
| chr17 | 42312484 | 42313455 | 972 | 4.66916 | 2.72445 | 1.90148 |
| chr17 | 42314875 | 42316383 | 1509 | 4.17741 | 2.58679 | 1.49223 |
| chr17 | 42337598 | 42338670 | 1073 | 4.51024 | 2.68161 | 1.76113 |
| chr17 | 42363913 | 42364381 | 469 | 3.57088 | 2.40576 | 1.01901 |
| chr17 | 42377801 | 42378844 | 1044 | 3.84965 | 2.41173 | 1.25401 |
| chr17 | 42387418 | 42389449 | 2032 | 4.75049 | 2.71867 | 1.98772 |

**Table S4.** Sample demographics of ChIP-seq donors and resulting data. Remaining reads count the number of trimmed, decontaminated reads following FASTQ preprocessing. The percentage of properly paired, aligned reads is given, as is the number of peaks identified by MACS2 for each sample relative to the input sample.

|  | **Sex** | **Age** | **Input Reads** | **Remaining Reads** | **% Properly aligned reads** | **# Enriched Peaks** |
| --- | --- | --- | --- | --- | --- | --- |
| **Input** | n/a | n/a | 156713964 | 156492276 | 99.27 | n/a |
| **p1035862-9** | F | 22 | 140187522 | 139974386 | 99.38 | 2412 |
| **p1035907-9** | F | 68 | 135521910 | 135246434 | 99.20 | 1012 |
| **p1035916-5** | F | 30 | 136026564 | 135808588 | 99.16 | 2181 |
| **p1035928-8** | F | 71 | 128434534 | 128086940 | 99.13 | 15210 |
| **p1035941-1** | M | 24 | 122478960 | 122245832 | 99.41 | 2866 |
| **p1035953-2** | M | 32 | 132110454 | 131866146 | 99.05 | 2756 |

**Table S5.** Significantly enriched functional categories for genes associated with ERα binding regions by GREAT. Significantly enriched categories met a binomial FDR Q-value threshold of 5%.

| **Ontology** | **# Term Name** | **Binom Rank** | **Binom FDR Q-Val** |
| --- | --- | --- | --- |
| GO Biological Process | DNA replication-dependent nucleosome assembly | 45 | 3.41115e-14 |
|  | protein refolding | 69 | 5.99735e-11 |
|  | positive T cell selection | 72 | 1.88179e-10 |
|  | positive regulation of nuclear-transcribed mRNA catabolic process, deadenylation-dependent decay | 79 | 3.96028e-10 |
|  | positive regulation of mRNA catabolic process | 80 | 4.38968e-10 |
|  | T-helper cell differentiation | 108 | 4.94657e-8 |
|  | positive regulation of natural killer cell activation | 113 | 1.59572e-7 |
|  | natural killer cell differentiation | 117 | 2.36310e-7 |
|  | chromatin silencing at rDNA | 120 | 2.89391e-7 |
|  | T-helper 17 cell differentiation | 168 | 1.13721e-5 |
|  | CD4-positive or CD8-positive, alpha-beta T cell lineage commitment | 197 | 6.09469e-5 |
|  | telomere capping | 209 | 7.80781e-5 |
|  | spliceosomal tri-snRNP complex assembly | 263 | 3.67336e-4 |
|  | stress granule assembly | 296 | 8.86802e-4 |
|  | ribosomal small subunit assembly | 355 | 3.47249e-3 |
| Cellular Component | nucleosome | 4 | 1.17346e-15 |
|  | DNA packaging complex | 8 | 4.23471e-13 |
|  | MHC protein complex | 9 | 8.12117e-13 |
|  | integral component of lumenal side of endoplasmic reticulum membrane | 21 | 5.32903e-6 |
|  | alpha-beta T cell receptor complex | 22 | 1.01614e-5 |
|  | MHC class II protein complex | 25 | 6.36248e-5 |
|  | PTW/PP1 phosphatase complex | 45 | 3.91793e-3 |
|  | MHC class I peptide loading complex | 49 | 7.12120e-3 |
|  | PRC1 complex | 53 | 1.44796e-2 |
| GO Molecular Function | peptide antigen binding | 9 | 8.11280e-9 |
|  | MHC class II protein complex binding | 26 | 5.08035e-5 |
|  | snRNA binding | 43 | 2.78861e-4 |
| Mouse Phenotype Single KO | abnormal memory T cell physiology | 23 | 7.41443e-16 |
|  | decreased B-1a cell number | 45 | 1.99471e-10 |
|  | abnormal thymocyte activation | 50 | 7.40996e-10 |
|  | decreased activated T cell number | 98 | 7.31205e-6 |
|  | decreased level of surface class I molecules | 100 | 1.08094e-5 |
|  | abnormal level of surface class I molecules | 107 | 2.37547e-5 |
|  | absent CD8-positive, alpha-beta T cells | 116 | 3.52385e-5 |
|  | abnormal antigen presentation via MHC class I | 153 | 2.74616e-4 |
|  | abnormal brachial lymph node morphology | 184 | 1.13170e-3 |
| Mouse Phenotype | abnormal memory T cell physiology | 11 | 3.96990e-23 |
|  | abnormal T cell selection | 17 | 2.49581e-15 |
|  | abnormal mature gamma-delta T cell morphology | 19 | 1.55604e-14 |
|  | abnormal positive T cell selection | 23 | 4.89923e-14 |
|  | decreased DN1 thymic pro-T cell number | 33 | 5.64557e-12 |
|  | abnormal thymocyte activation | 46 | 6.33874e-10 |
|  | abnormal gamma-delta intraepithelial T cell morphology | 48 | 1.27486e-9 |
|  | decreased spleen germinal center size | 49 | 1.28510e-9 |
|  | decreased level of surface class I molecules | 82 | 6.58477e-7 |
|  | abnormal level of surface class I molecules | 88 | 1.84828e-6 |
|  | absent B-1a cells | 89 | 2.44495e-6 |
|  | absent B-1 B cells | 92 | 3.72019e-6 |
|  | abnormal antigen presentation via MHC class I | 107 | 3.32432e-5 |
|  | decreased lymphoma incidence | 115 | 4.65799e-5 |
|  | abnormal IgG2c level | 184 | 1.21934e-3 |
|  | decreased activated T cell number | 244 | 9.42092e-3 |

**Supplemental Figures**

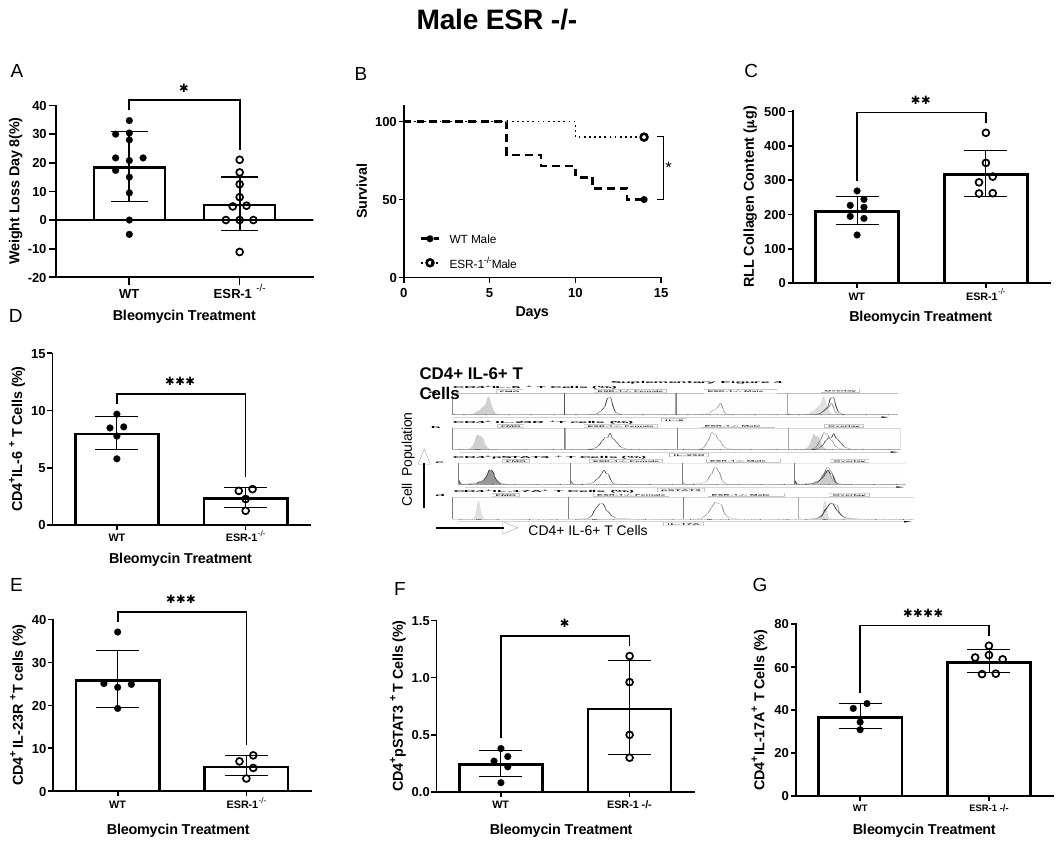

**Supplemental Figure 1: Loss of estrogen receptor alpha subunit (ESR-1) improves male survival and augments fibrosis. A)** WT and ESR-/- mice were treated with bleomycin and monitored for 14 days A) Body weights of mice at day 8 compared to day 0; **B)** Murine mortality across 14 days. Kaplan-Meier survival analysis with log-rank test was used to determine differences between groups; **C)** Soluble collagen content of RLL by Sircol assay. Flow cytometric analysis of T cells from single cell lung suspensions at day 14 for **D)** IL-6; **E)** IL-23R; **F)** pSTAT3Y^705^ and **G)** IL-17A. Comparisons between cohorts were performed using one-way ANOVA with Tukey’s post-hoc. *P < 0.05, **P < 0.01, ***P < 0.001, ****P < 0.0001, ns: no significance. RLL: Right lower lobe. Bars are mean ± SD; each symbol represents an individual mouse. WT: Wildtype, ESR-1-/-: estrogen receptor alpha knockout mice.

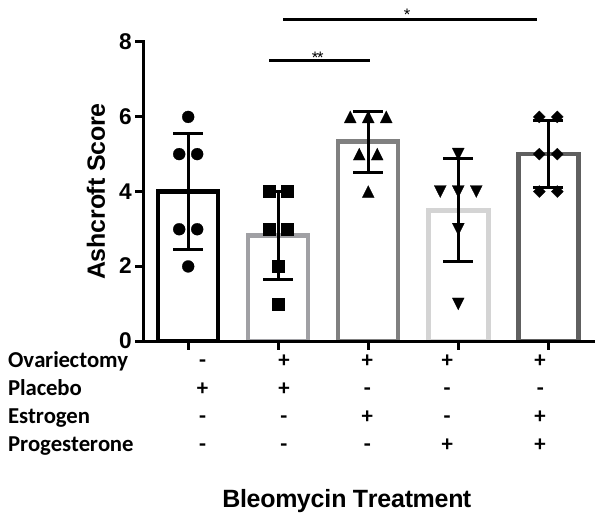

**Supplemental Figure 2. Ovariectomized mice display decreased lung fibrosis.** Ashcroft scoring of sham and ovariectomized mice treated with placebo, 17β-E2, progesterone, or both. Comparisons between cohorts were performed using student’s one-way ANOVA *P < 0.05, **P< 0.01
